# Unique Protein Interaction Networks Define The Chromatin Remodeling Module of The NuRD Complex

**DOI:** 10.1101/2021.01.27.428018

**Authors:** Mehdi Sharifi Tabar, Caroline Giardina, Yue Julie Feng, Habib Francis, Hakimeh Moghaddas Sani, Jason K. K. Low, Joel P. Mackay, Charles G. Bailey, John E.J. Rasko

## Abstract

The combination of four proteins and their paralogues including MBD2/3, GATAD2A/B, CDK2AP1, and CHD3/4/5, which we refer to as the MGCC module, form the chromatin remodeling module of the Nucleosome Remodeling and Deacetylase (NuRD) complex, a gene repressor complex. Specific paralogues of the MGCC subunits such as MBD2 and CHD4 are amongst the key repressors of adult-stage fetal globin and provide important targets for molecular therapies in beta (β)-thalassemia. However, mechanisms by which the MGCC module acquires paralogue-specific function and specificity have not been addressed to date. Understanding the protein-protein interaction (PPI) network of the MGCC subunits is essential in defining underlying mechanisms and developing treatment strategies. Therefore, using pulldown followed by mass spectrometry analysis (PD-MS) we report a proteome-wide interaction network of the MGCC module in a paralogue-specific manner. Our data also demonstrate that the disordered C-terminal region of CHD3/4/5 is a gateway to incorporate remodeling activity into both the ChAHP (CHD4, ADNP, HP1γ) and NuRD complexes in a mutually exclusive manner. We define a short aggregation prone region (APR) within the C-terminal segment of GATAD2B that is essential for the interaction of CHD4 and CDK2AP1 with the NuRD complex. Finally, we also report an association of CDK2AP1 with the Nuclear Receptor Co-Repressor (NCOR) complex. Overall, this study provides insight into the possible mechanisms through which the MGCC module can achieve specificity and diverse biological functions.

## Introduction

The NuRD complex plays a crucial function in embryonic stem cell biology and development, and cancer [1-3]. The NuRD complex consists of seven core subunits, namely MTA, HDAC, RBBP, MBD, GATAD2, CDK2AP1, and CHD, with each subunit interchangeable with its paralogous family members: MTA1, MTA2 and MTA3; HDAC1 and HDAC2; RBBP4 and RBBP7; MBD3 and MBD2; GATAD2A and GATAD2B; CHD3, CHD4 and CHD5 [4-7]. This ~1 MDa nuclear protein complex harbours both chromatin remodeling and lysine deacetylase activities [7, 8]. A symmetric module encompassing MTA, HDAC and RBBP proteins (MHR) conveys the lysine deacetylation activity via the HDAC catalytic subunit [8-11], whereas MBD, GATAD2, CDK2AP1, and CHD proteins (MGCC) collaborate to translocate nucleosomal DNA via the CHD subunit [12,13]. The canonical NuRD complex comprises a stable asymmetric composition of two MHR modules and a single MGCC module [11]. Among other functions, MGCC components regulate the expression of adult globin genes and are linked to *β*-hemoglobinopathies [14-16]. Therefore, subunits of MGCC are considered as potential molecular targets for therapeutic activation of fetal gamma globin genes [17]. Understanding the auxiliary binding partners of the MGCC module will facilitate understanding of the molecular basis of MGCC-mediated gene silencing.

The domains responsible for mediating intra-NuRD architecture are being uncovered through careful biochemical analysis. Previous structural studies have documented that two HDAC molecules are activated by homo- or heterodimerisation of two molecules of MTA through ELM2-SANT domains [18]. MTA1/2, and MTA3 provide two and one binding regions, respectively, within their C-terminus for RBBP proteins to facilitate their access to histone tails to possibly remove acetyl marks from lysine residues [9,8]. The bromo-adjacent homology (BAH) domain of MTA proteins is well conserved but is structurally and functionally poorly characterised [19]. MBD connects to the N-terminal region of GATAD2 protein through a coiled-coil interface [20]. GATAD2 uses the GATA zinc finger domain in the C-terminal region to facilitate CHD binding [15,7].

Focusing on the MGCC module, MBD2 and MBD3, which are highly homologous and mutually exclusive within the NuRD complex [21], structurally bridge GATAD2/CDK2AP1/CHD subunits with the MHR module to form the intact NuRD complex [10,11]. Although MBD2 and MBD3 are highly similar, their affinity and selectivity toward DNA is markedly different [22,23]. MBD3 has been extensively studied but the function of MBD2 isoforms MBD2a and MBD2b are poorly understood. MBD2a contains an extra 148 amino acid Gly- and Arg-rich domain at the N-terminal end as compared to MBD2b [24,25]. Within GATAD2, the C-terminal portion is responsible for the interaction with CHD proteins [15,7]. However, the minimal region mediating this interaction has remained elusive. To date, CDK2AP1 is the least studied subunit of NuRD biochemically and possibly overlooked in some studies because of its small size. CHD4 contains several functional domains essential for ATP-dependent nucleosome remodeling. The N-terminal region of CHD4 binds DNA via an HMG-box-like domain [26]. CHD4 can bind to histone tails (such as H3K4 and trimethylated H3K9) via two plant homeodomain (PHD) zinc fingers and remodels nucleosomes via its ATPase motor domain assisted by tandem chromodomains (CHD) [27]. The C-terminal region of CHD4, however, is functionally and structurally the least characterised part of CHD4.

Recently, the NuRD subunit CHD4 was identified as a component of the ChAHP complex, which also comprises ADNP and HP1γ (otherwise known as CBX3) subunits [28]. ChAHP modulates chromatin organisation by neutralising loops formed by CTCF at specific genomic regions and plays an important role in embryonic neural development and heterochromatin organisation [29]. ADNP is the requisite subunit, without which the ChAHP complex cannot form [28]. How CHD4 can be independently involved in two different chromatin binding complexes, and whether CHD3 and CHD5 can also form stable complexes with ADNP has not been resolved. The minimal domains for these interactions are also unknown.

Based on the above understanding, we set out to address the following: (i) what proteins interact with MBD2 isoforms; (ii) what is the minimal region in GATAD2 proteins that mediates its interaction with CHD family members; (iii) is CDK2AP1 an exclusive subunit of NuRD or is it found in other complexes; and (iv) what regions in CHD4 and ADNP mediate their interaction? We have sought to address these questions by focusing on the interaction network of the MGCC module of NuRD and core component of the ChAHP complex, ADNP. We define novel factors and complexes that can interact with paralogues of the NuRD subunits. We also refine minimal interacting domains between certain subunits and isoforms of the NuRD MGCC module. The importance of these results lies in providing clarification of how the MGCC module functions to co-ordinate diverse cellular functions.

## Results

The domain architecture of the NuRD and ChAHP complexes is depicted showing known domains and sites of interaction **(Fig. 1A)**. Domains responsible for mediating intra-NuRD subunit interactions that have not previously been described or characterised are highlighted with a question mark **(Fig. 1A)**.

**Fig. 1.**
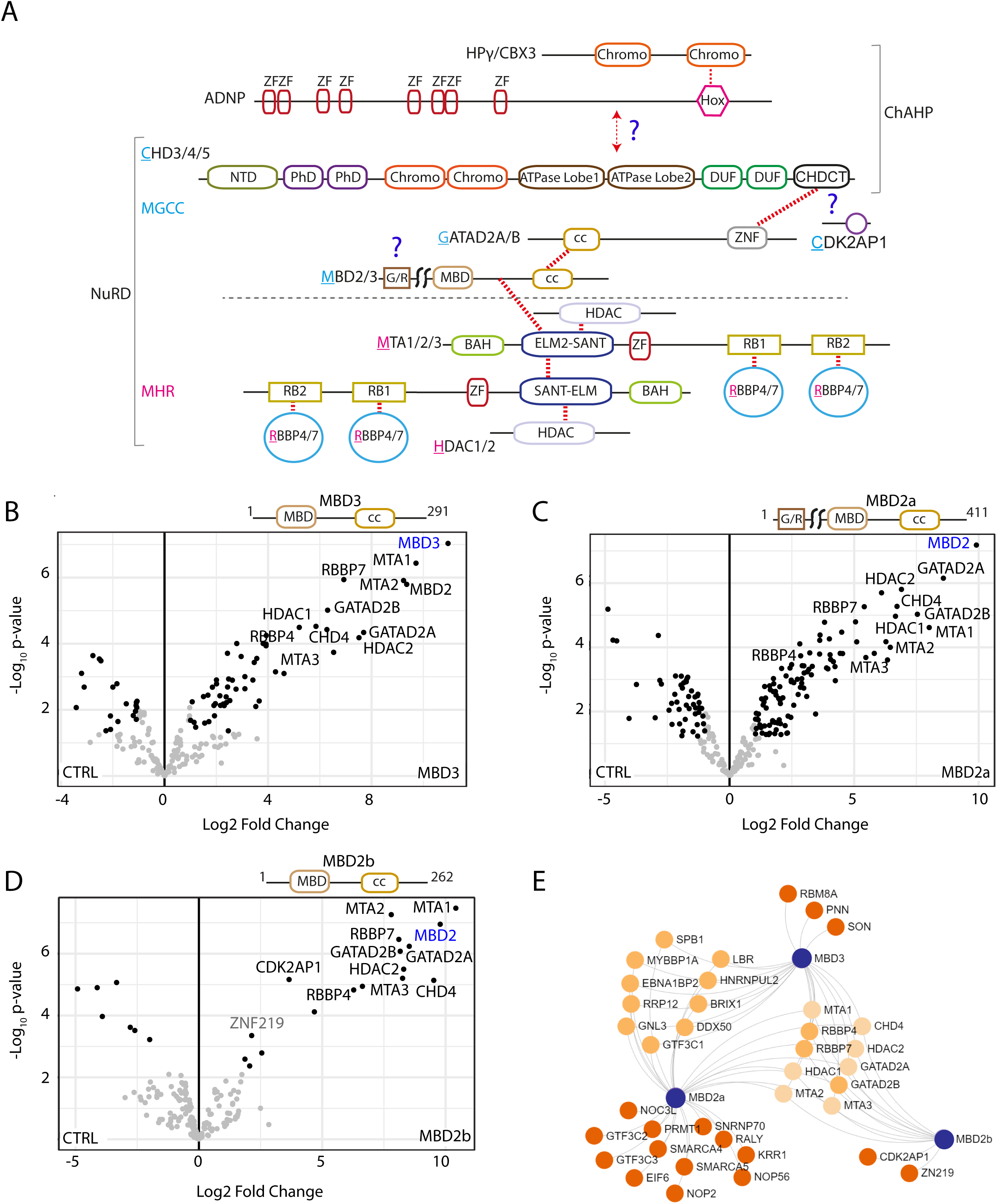
Mutual and distinct MBD binding partners revealed by PD-MS. **A)** Schematic diagram depicting the domain organization of the NuRD and ChAHP complexes. Coloured regions indicate domains with known structures, or which are predicted to be ordered; and single lines specify regions predicted to be disordered. The total NuRD complex contains MHR and MGCC modules separated by a dashed line. CHD4 is a component of both ChAHP and NuRD complexes. Red dotted lines represent the inter-subunit binding regions. Question marks indicate the regions of interaction that have not been well defined. **B-D)** FLAG pulldowns were performed in triplicate followed by label-free quantitative mass spectrometry analysis and LFQ intensities were used to generate the volcano plots. The canonical subunits of the NuRD complex are highlighted in black, and non-canonical or potential new interactors in grey. Significantly enriched proteins in **B)** MBD3 **C)** MBD2a and **D)** MBD2b pulldowns. Significantly enriched proteins are indicated with black dots; non-significant proteins are indicated with grey dots; the bait is indicated as blue text. **E)** Network representation of shared and specific interactors of MBD proteins. Significantly enriched proteins with at least 2-fold change were used to generate the network. A full list of interactors is available in **Supplemental Table 1**.

### MBD3 and MBD2 isoforms have both mutual and unique binding partners

Initially, we focused on the MBD subunits, which bridge the MGCC to the MHR module. Relative <u>L</u>abel-<u>F</u>ree <u>Q</u>uantification (LFQ) analysis of MBD2a, MBD2b and MBD3 PD-MS data demonstrate that all three MBDs significantly co-purify the canonical subunits of the NuRD complex (**Fig. 1B-D, Supplemental Table 1, S1 & S2**). Both MBD2a and MBD3 mutually co-immunoprecipitated a substantial number of possible new interactors involved in chromatin biology, transcriptional gene regulation, and RNA processing (**Fig. 1B & C**). MBD2b had comparatively few interactors, but significantly enriched the canonical NuRD subunit CDK2AP1 (**Fig. 1D**). Interestingly, CDK2AP1 was significantly enriched only with the MBD2b isoform but not MBD2a, or MBD3. This shows the preference of CDK2AP1 for binding to the MBD2/NuRD complex but not the MBD3/NuRD **(Supplemental Table 1, S1**). We also identified other proteins specifically enriched with each MBD family member. For example, the MBD2a isoform, which contains an N-terminal Gly- and Arg-rich domain, captures numerous proteins, including the chromatin remodeling proteins SMARCA4 and SMARCA5 as well as the arginine N-methyltransferase enzyme PRMT1 (**Fig. 1E**). Evidence for arginine methylation of MBD2a [30] suggests a possible mechanism by which the function of the MBD2/NuRD complex is regulated. Interestingly, ZNF219, a transcriptional repressor that is a known interactor of the NuRD complex, was only pulled down with MBD2b (**Fig. 1D, E** and **Supplemental Table 1, S1** & **S2**) [6]. Given that MBD2b is not the major isoform of MBD2, this supports the sub-stoichiometric nature of ZNF219 noted previously [6] and possibly other non-canonical partners of the NuRD complex that have been observed. Our data reveal that the two MBD2 isoforms show distinct molecular interactions and could form NuRD complexes associating with distinct sets of partner proteins. Hence, these diverse NuRD sub-species might perform different gene regulatory activities at different loci.

### A 40-residue region in the GATAD2B C-terminus is important for connecting CHD4 to the NuRD complex

Previous biochemical studies have demonstrated that the C-terminal region (residues 276-593) in GATAD2 proteins is necessary for binding to CHD4 [15,7]. Using GATAD2B as an exemplar, we examined whether other proteins can compete with CHD4 for binding to GATAD2B and then defined the minimal region that mediates this interaction. LFQ analysis of proteins co-purified with full-length GATAD2B showed marked enrichment of the canonical NuRD subunits (**Fig. 2A**). We also observed significant enrichment of auxiliary proteins important for chromatin biology and gene repression, namely BEND3 and KCRM. PD-MS of the N-terminal half of GATAD2B (GATAD2B-N, residues 1-276) revealed that all canonical NuRD subunits except for CDK2AP1 and CHD4 were enriched (**Fig. 2B** and **Supplemental Table 1, S1** & **S3**).

**Fig. 2.**
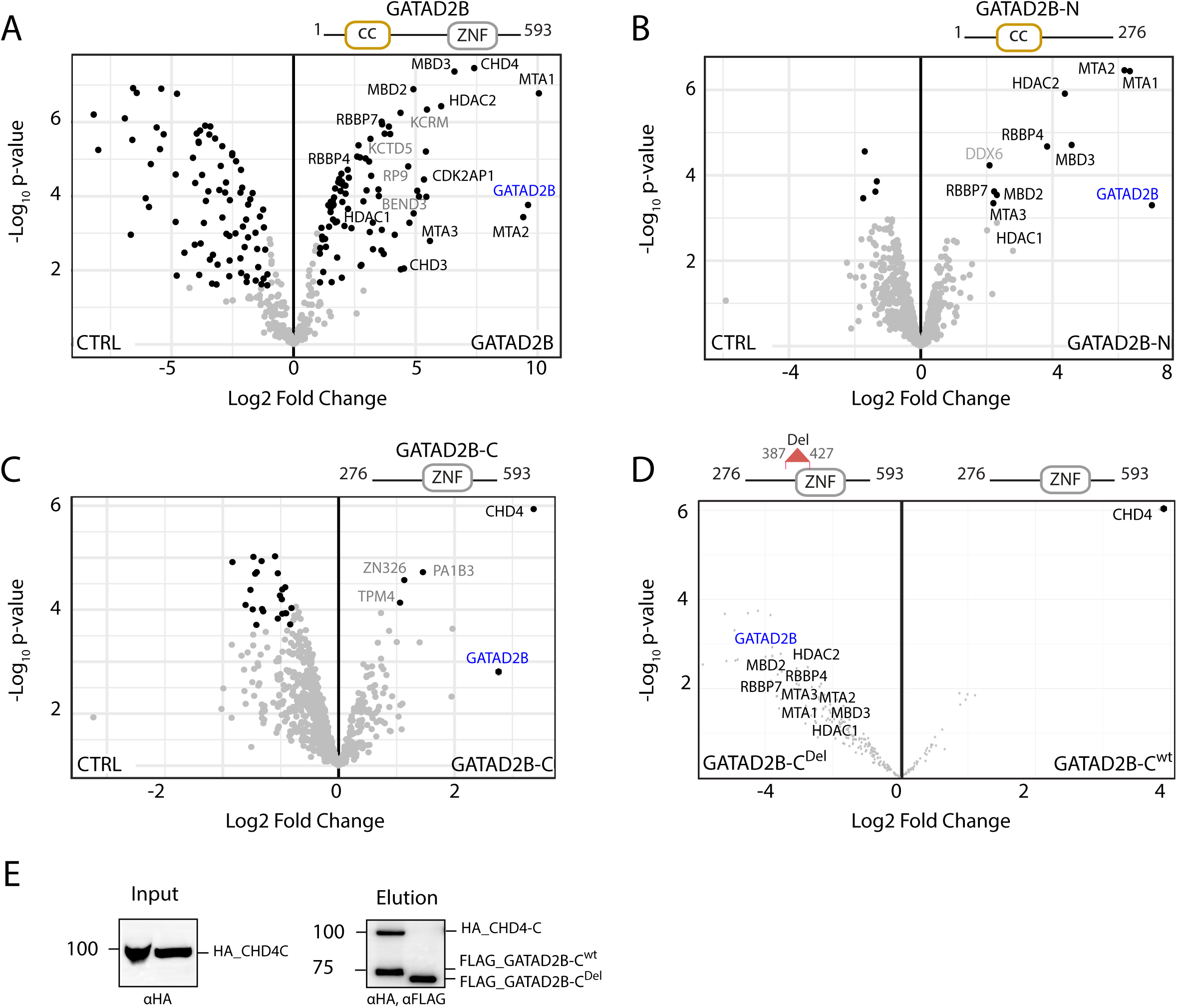
GATAD2B C-terminus mediates interaction with CHD4. Volcano plots were generated using LFQ intensities of FLAG PD-MS and the canonical subunits of the NuRD complex are highlighted in black, and non-canonical or potential new interactors in grey. **A)** Full-length GATAD2B **B)** N-terminus (GATAD2B-N) and **C)** C-terminus (GATAD2B-C). FLAG-only was used as a control (CTRL) to account for background contamination. **D)** FLAG PD-MS of GATAD2B-C^Del^ versus WT GATAD2B-C. Schematic of GATAD2B constructs used as baits are indicated on top of the volcano plots. **E)** Western blots of GATAD2B proteins were co-expressed in the IVT system and purified on anti-FLAG beads. Input (5% of total, left panel) blot was developed with anti-HA and the Elution (70% of total, right panel) was developed with both HA and FLAG antibodies.

On the other hand, using GATAD2B C-terminal region (GATAD2B-C) as bait, CHD4 was the only NuRD subunit that prefers the C-terminal portion of GATAD2B for binding (**Fig. 2C**); it was also the most highly enriched protein in this experiment across the whole proteome. Examination of iBAQ (<u>i</u>ntensity-<u>B</u>ased <u>A</u>bsolute <u>Q</u>uantification) values in all GATAD2B bait replicates revealed that all three replicates of GATAD2B (full-length) and two replicates of GATAD2B-C but not GATAD2B-N had high intensity values for CDK2AP1 (**Supplemental Fig. 1** and **Supplemental Table 1, S1**), indicating the C-terminal GATAD2B binding preference of CDK2AP1.

To narrow down the CHD4 binding region in the C-terminus of GATAD2B, residues 387-427 of GATAD2B were deleted from GATAD2B-C (herein named GATAD2B-C^Del^) and PD-MS was performed. This pulldown showed marked depletion of CHD4 (but not other NuRD subunits), compared to the wild-type GATAD2B-C (**Fig. 2D** and **Supplemental Table 1, S1**). Notably, this deletion did not disrupt the interaction of other NuRD subunits. To corroborate this finding, we performed co-immunoprecipitation using FLAG-tagged GATAD2B-C or GATAD2B-C^Del^ versus an HA-tagged C-terminal CHD4 construct (HA-CHD4-C, residues 1230-1912) using an *in vitro* translation (IVT) system. IVT confirmed that the GATAD2B-CHD4 interaction was direct and that residues 387-427 are necessary for the integrity of this interaction (**Fig. 2E**).

### The GATAD2 interaction interface with CHD4 includes an aggregation prone region (APR)

Protein-protein interaction interfaces typically contain high proportions of hydrophobic residues. These interfaces can display characteristics of aggregation prone regions with fibril forming capacities. Analysis of GATAD2A and GATAD2B protein sequences using the TANGO aggregation prediction algorithm [31], revealed a number of mutually exclusive as well as overlapping APRs (**Fig. 3A**). The most prominent of these APRs was a small and highly similar stretch of seven hydrophobic and aliphatic amino acids in GATAD2A (residues 384-390, FIYLVGL) and GATAD2B (residues 388-394 – FIYMVGL) (**Fig. 3A & B**). We designed peptidomimetics containing the 7-residue APR regions from GATAD2A and GATAD2B conjugated to a 11-residue portion of HIV-1 Tat protein for cell permeability, referred to herein as APR^A^ and APR^B^, respectively. We first tested the impact of these APR peptides on K562 cell viability compared to a Tat-conjugated control peptide comprising seven alanine residues (CTRL). After dose-dependent addition of APR^A^, APR^B^ and CTRL peptides K562 cell proliferation was analysed after 48 h by MTT (3-(4,5-dimethylthiazol-2-yl)-2,5-diphenyltetrazolium bromide) assay. APR^A^ and APR^B^ induced cell death with IC50 values of ~30 ΜM whereas the CTRL peptide had no effect (**Fig. 3C**).

**Fig. 3.**
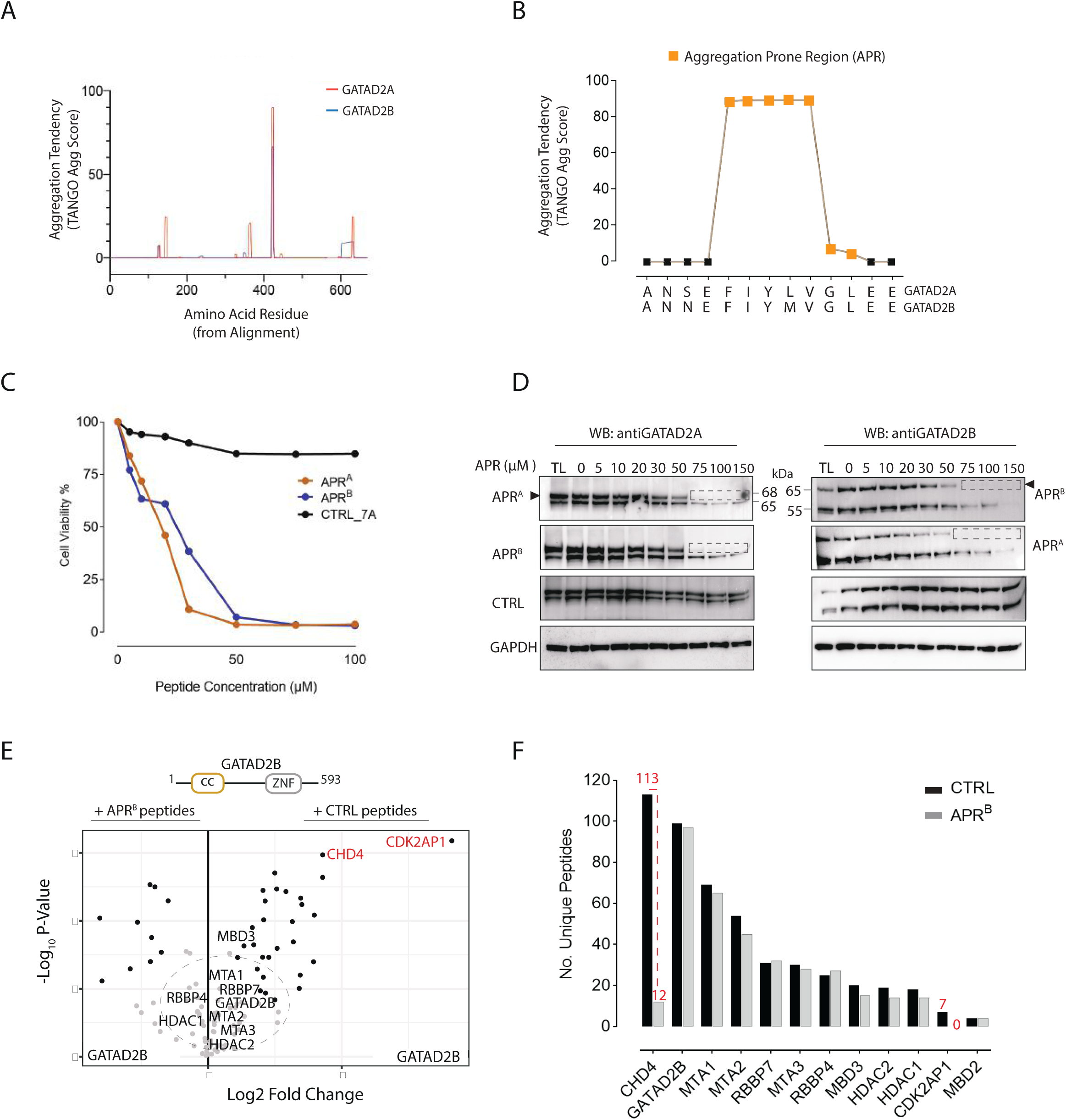
An aggregation prone region within GATAD2 proteins co-ordinates CDK2AP1 and CHD4 association with NuRD. **A)** TANGO analysis show β-sheet aggregation tendency for the APR region within GATAD2A and GATAD2B **B)** A close-up schematic of the APRs and gatekeeper residues (388-394) in GATAD2 proteins. **C)** Graph representing the MTT assay performed in 2 biological and three technical replicates (n=6). Cell viability of K562 cells treated with a range of APR peptide concentrations was measured after 48 h. Data are presented as a percent of untreated cell viability measured using the MTT assay. **D)** Western blots of endogenous GATAD2 proteins after treatment of K562 cell lysates with APR-tat peptides. APR peptidomimetics (0-150 μM) were added to the lysates before sonication. Total lysates (TL, 5%) were taken as input before separation of the soluble and insoluble fractions by centrifugation. Cleared lysate (5% of soluble fraction) was run on SDS-PAGE and probed for GATAD proteins. Black arrows indicate the canonical isoform of GATAD2A and GATAD2B proteins, and dotted rectangular box shows loss of soluble proteins post APR treatment. **E)** Volcano plot comparing the interactors of over-expressed GATAD2B in the presence of 50 μM APR^B^ (left panel) or CTRL (right panel) peptides. **F)** The number of unique peptides of each subunit of the NuRD complex detected by mass spectrometry after FLAG-GATAD2B PD-MS in the presence of APR^B^ and CTRL peptides.

To confirm whether these peptidomimetics were inducing protein aggregation, we titrated APR peptidomimetics into K562 total cell lysates (after sonication and before centrifugation). Addition of APR^A^ and APR^B^ resulted in the aggregation and depletion of soluble endogenous GATAD2A and GATAD2B proteins whereas the control peptidomimetic had no impact on GATAD2 protein solubility (**Fig. 3D**). Due to the similarity between the GATAD2A/B APR regions (**Fig. 3B**), APR^A^ and APR^B^ depleted GATAD2B and GATAD2A equally, indicating no specificity for a particular GATAD2 protein (**Fig. 3D**). We next performed FLAG-GATAD2B PD-MS in the presence of 50 µM APR^B^ or CTRL to determine if the APR mimetic peptides could directly interfere with any PPIs that require the APR region (**Fig. 3E, Supplemental Table1, S4**). We observed a significant loss of NuRD subunits CDK2AP1 and CHD4 in the presence of APR^B^ peptides compared to the CTRL, while no significant change was observed for GATAD2B and other NuRD subunits (**Fig. 3E** and **Supplemental Table 1, S4**). We also examined the total number of peptides detected for all NuRD subunits to ensure that the loss of CHD4 and CDK2AP1 was not due to a reduction in GATAD2B or other subunits of the NuRD complex. Interestingly, we saw no significant difference in the total number of unique peptides (99 vs 97) of GATAD2B and NuRD subunits other than CHD4 and CDK2AP1 (**Fig. 3F** and **Supplemental Table 1, S4**). In contrast, the number of unique peptides for CHD4 and CDK2AP1 decreased from 113 to 12 and 7 to 0, respectively (**Fig. 3F**). Based on these results, we conclude that APR^B^ competes away CHD4 and CDK2AP1 from binding to the NuRD complex.

### CDK2AP1 interacts with both the NuRD and NCOR complexes

CDK2AP1 is a 12-kDa protein involved in cell cycle regulation and is the least defined subunit of the NuRD complex. Pulldown of a CDK2AP1-GFP fusion previously demonstrated that CDK2AP1 is a canonical component of the NuRD complex [32]. To corroborate these findings, we used the smaller (∼1 kDa) FLAG tag to immunoprecipitate CDK2AP1, and our PD-MS confirmed the enrichment of canonical NuRD subunits using medium stringency washes (**Fig. 4A**). With high stringency washes (500 mM NaCl), the interaction of CDK2AP1 with CHD4 and GATAD2 proteins in particular was not abrogated, suggesting this is a high affinity and direct interaction (**Fig. 4C, D** and **Supplemental Table 1, S5**).

**Fig. 4.**
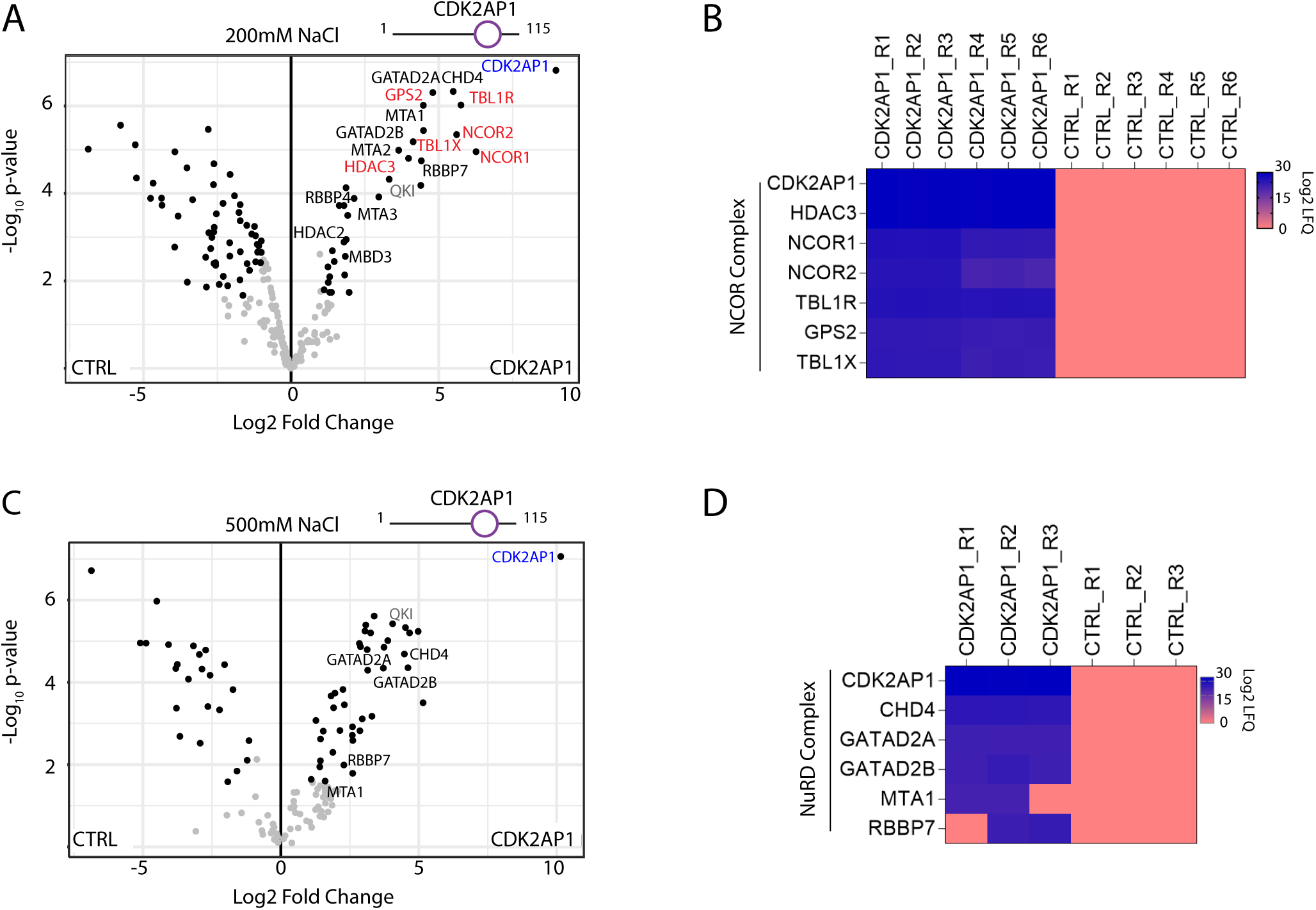
CDK2AP1 interacts with NuRD and NCOR complexes. **A)** & **C)** Volcano plots represent enrichment of the NuRD and NCOR subunits in FLAG-CDK2AP1 and FLAG-only CTRL pulldowns in the presence of: A) 200 mM NaCl (n=6); and C) 500 mM NaCl (n=3). **B)** & **D)** LFQ intensity-based heatmap of FLAG-CDK2AP1 PD-MS versus FLAG alone control (CTRL) showing: **B)** NCOR complex subunits pulled down only with CDK2AP1 and not in any replicates of CTRL in presence of 200 mM NaCl; **D)** Canonical NuRD subunits at 500 mM NaCl.

Notably, we also observed enrichment of NCOR1, NCOR2, TBL1X, TBL1R, GSP2, and HDAC3, which are all canonical subunits of the NCOR complex (**Fig. 4A**). An LFQ intensity-based heatmap of the NCOR subunits detected in FLAG-CDK2AP1 versus CTRL samples confirmed that the interaction was not mediated by beads or FLAG tag (**Fig. 4B, Supplemental Table 1, S5**). This result is supported by the number of unique NCOR complex peptides obtained as well as the low representation of these proteins in databases of common mass spectrometry contaminants (**Supplemental Table 2**). Whether CDK2AP1 is a novel integral subunit or only an interactor of the NCOR complex was not resolved. However, at 500 mM NaCl, all interactions of CDK2AP1 with the NCOR complex subunits were lost, suggesting that this interaction might be of low to medium affinity (**Fig. 4C**). Interestingly, the RNA-binding protein QKI, which is involved in mRNA stability, translation, and splicing was also significantly enriched in CDK2AP1 PD-MS, even with stringent washing conditions (**Fig. 4A & C**). CDK2AP1-QKI is a strong and possibly direct interaction but occurs independently of the MGCC module as it was not detected in other PD-MS experiments in this study.

### The C-terminal end of CHD proteins facilitates engagement with the NuRD and ChAHP complexes

Our previous *in vitro* co-IP studies of NuRD subunits revealed that the CHD4 C-terminus (aa 1230-1912) interacts with GATAD2 proteins [7]. In addition, a recent report showed that CHD4 also interacts with the ChAHP complex [15]. Given that the NuRD complex can also incorporate CHD3 or CHD5, we investigated the interactome networks mediated through the C-termini of all NuRD-associated CHD proteins. Accordingly, FLAG-tagged CHD3-C, CHD4-C and CHD5-C complexes were purified and analysed by mass spectrometry. We observed strong enrichment of most NuRD canonical subunits (**Fig. 5A, B & C**). Interestingly, CDK2AP1 enrichment was seen only with CHD4-C but not with CHD4-N and CHD4-M (**Fig. 5D & Supplemental Fig. 2)**, indicating that CDK2AP1 only recognizes the C-terminal half of CHD4 (**Fig. 5A**).

**Fig. 5.**
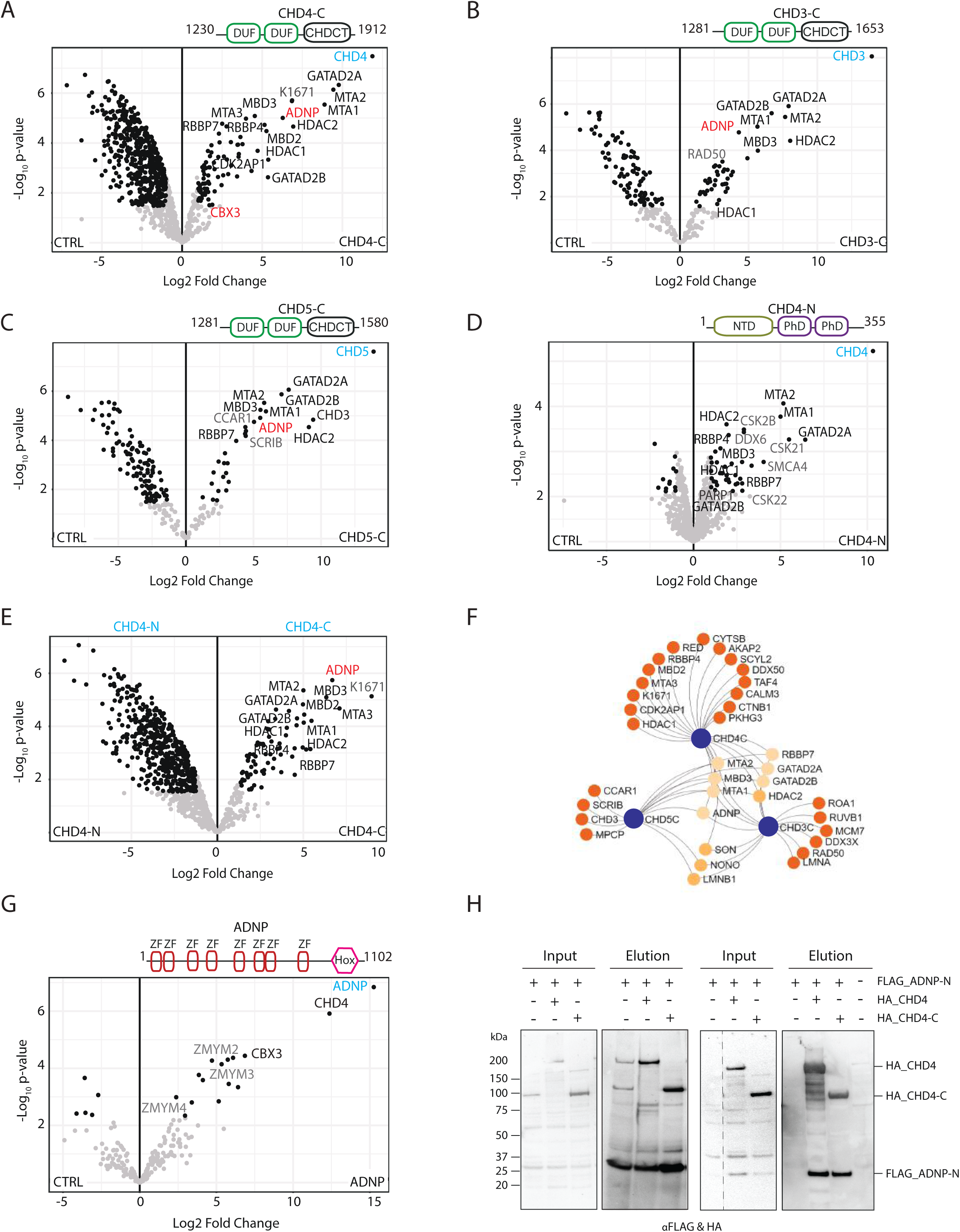
CHD3/4/5 C-termini preferentially bind NuRD subunits and ADNP. The canonical subunits of the NuRD are highlighted in black, ADNP in red and potential new interactors in grey. Interaction partners of **A)** CHD4-N **B)** CHD4-C **C)** CHD3-C and **D)** CHD5-C compared to CTRL, and **E)** CHD4-N vs CHD4-C **F)** Network representation of shared and specific interactors of CHD proteins. Significantly enriched proteins with at least 2-fold change were used to generate the network. Bait proteins are indicated in blue circles, and unique interactors of each family member are indicated in dark orange and shared interactors in light orange. **G)** ADNP co-purifies with CHD4 and CBX3/HP1γ, known subunits of ChAHP complex. ZMYM family proteins are indicated in grey. A full list of the interactions is available in Supplemental Table 1. **H)** Western blots of input and elution samples from Flag-ADNP-N (1-228 aa) co-expressed with CHD4 protein. Left, pulldown showing that FLAG-ADNP-N purified on anti-FLAG beads pulls down co-expressed CHD4 full-length (Lane2) or CHD4-C (Lane 3). Right, pulldown showing that CHD4 full-length (Lane2) or CHD4-C (Lane 3) purified on anti-HA beads pulls down co-expressed FLAG-ADNP-N.

PD-MS performed with the CHD4 N-terminal region (CHD4-N) also showed enrichment of NuRD subunits, suggesting that CHD4-C is not the sole region responsible for engagement with NuRD (**Fig. 5D**). However, notable enrichment of NuRD subunits was seen whit CHD4-C when compared to CHD4-N (**Fig. 5E**). Notably, we observed enrichment of CSK2B (CSNK2B), CSK21 (CSNK2A1) and CSK22 (CSNK2A2), well-known subunits of the casein kinase II (CSK2) serine/threonine protein kinase complex, in the CHD4-N pulldown (**Fig. 5D**). The CSK2 complex regulates the function of many regulators of chromatin organisation and function and has been shown to phosphorylate serine residues in the CHD4 N-terminus [33]. We also observed specific enrichment of activity-dependent neuroprotective protein (ADNP) with all CHD family members (**Fig. 5A-C, F** and **Supplemental Table 1, S6)**. Since the discovery of the ChAHP complex containing ADNP, CBX3 and CHD4 in 2017 [28], attention has focused on its molecular and cellular function in cancer and neurodegenerative diseases. It has been shown by PD-MS of full-length FLAG-tagged ADNP in HEK293 cells that CHD4 and CBX3 are the top-enriched proteins [28]. In addition, we saw marked enrichment of three members of the MYM-type zinc finger family, namely ZMYM2, ZMYM3 and ZMYM4 (**Fig. 5G**, and **Supplemental Table 1, S6**). Significant enrichment of these proteins with ChAHP may suggest a role for MYM-type proteins in genome organisation and architecture. To corroborate CHD4 interaction with ADNP, we performed co-immunoprecipitation using FLAG-tagged ADNP N-terminus (ADNP-N, aa 1-228) and full-length HA-CHD4 or HA-CHD4-C (aa 1230-1912) proteins in HEK293 cells. FLAG-ADNP-N immobilised on FLAG beads could pulldown both full length HA-CHD4 and the HA-CHD4-C (**Fig. 5 H**, left panel). Similarly, HA-CHD4 proteins immobilised on HA beads could also precipitate FLAG-ADNP-N (**Fig. 5H**, right panel). Notably, the C-terminal half of ADNP failed to express (data not shown).

### Cancer missense mutations may change the balance of CHD/NuRD and CHD/ChAHP complexes

The impact of somatic or germline mutations on large, multi-subunit complexes such as NuRD is being recognised. We focused on CHD4 as a mutually exclusive partner of both NuRD and ChAHP complexes to determine whether any cancer-specific missense mutations in the C-terminal region of CHD4 disrupted its binding to either GATAD2 or ADNP proteins. Previous CRISPR/Cas9 screens in erythroid cells revealed that disruption of aa 1872-1883 of CHD4 abrogated the CHD4-GATAD2B interaction [15]. We therefore examined the six missense mutations in CHD4 within or adjacent to this region, which could potentially affect the interaction of CHD4 with either ADNP or GATAD2B **(Supplemental Fig. 3A)**. To understand the effect of the CHD4 C-terminal missense mutations in the context of NuRD and ChAHP assembly, we performed pairwise interaction experiments to evaluate the binding of CHD4-C to full-length ADNP and GATAD2B proteins. Wild type (WT) or mutant HA-tagged CHD4-C as well as FLAG-tagged ADNP and GATAD2B proteins were co-expressed and their interaction examined in pulldowns followed by western blot experiments. Four of these mutations (D1867N, P1879S, R1890C, and N1891D) had no impact on CHD4 interaction with either GATAD2B or ADNP (D1867N is exemplified in **Supplemental Fig. 3B**, Lane 2). However, a reduction in CHD4-C^A1866D^ interaction with full length ADNP was observed, when compared to WT CHD4-C **(Supplemental Fig. 3B)**. With regard to GATAD2B, only CHD4-C^E1889K^ showed a clear effect on the interaction when compared to WT **(Supplemental Fig. 3C)**. These mutations were not completely disruptive but reduced the affinity of CHD4 for either NuRD or ChAHP subunits. These results may suggest that the composition and proportion of CHD4-NuRD and CHD4-ChAHP complexes might be perturbed in cancer cells carrying these particular CHD4-C mutations.

### Fixed and altered stoichiometries was observed for MHR and MGCC modules, respectively

Understanding the stoichiometry of the NuRD complex subunits will help delineate its structure and function. To this aim, we used the iBAQ-adjusted intensity values as described previously [34,6]. Because of the high sequence similarity between paralogues (i.e., RBBP4 is 92% identical to RBBP7), we first considered each set of paralogues as a single group and averaged their iBAQ values to assess stoichiometry. The averaged values were divided by the averaged MTA value and multiplied by 2 because based on the published X-ray crystal structure two molecules of MTA are found in an intact NuRD complex (**Supplemental Table 1, S7**). Of note, bait proteins were excluded because they are in excess and introduce a bias in stoichiometry calculations. MBD and GATAD2 have been considered as exclusive subunits of the NuRD complex. Thus, we used their iBAQ data for stoichiometry calculations. Using MBD PD-MS data we calculated stoichiometric ratios of ~1:0.3:0.1:2:2:4 for GATAD2:CHD:CDK2AP1:MTA:HDAC:RBBP4 (**Fig. 6A**). The 2:2:4 ratios for MHR exactly matches the previous crystal and NMR structures [18], as well as MS quantifications [35,11,6,34]. Similarly, when GATAD2B was used as bait, the ratio for the MHR module remained constant (2:2:4) but we calculated a ratio of ∼2:1:2 for MBD:CHD:CDK2AP1 (**Fig. 6B**).

**Fig. 6.**
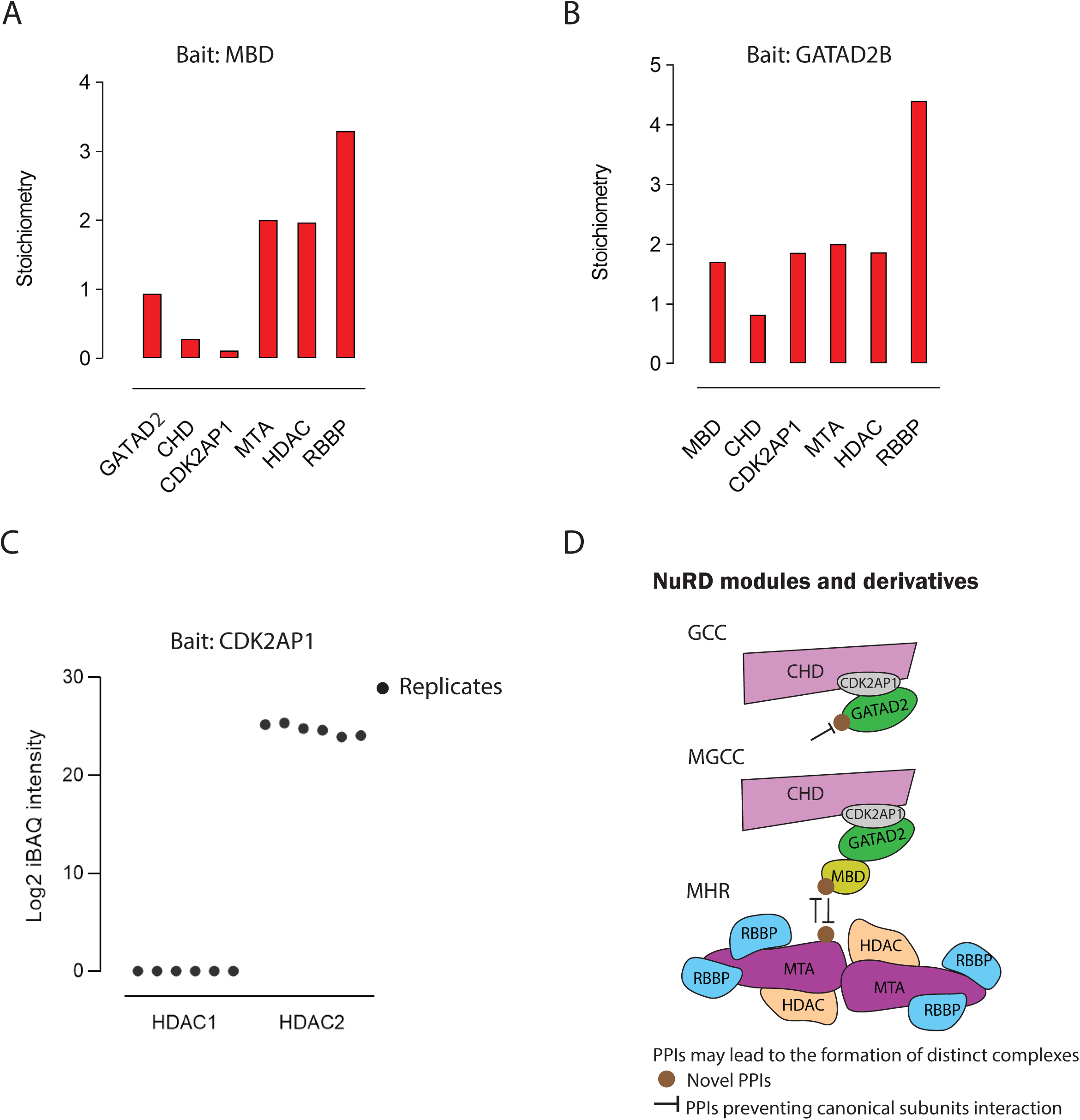
Stoichiometry of the NuRD subunits. MBD2/3 and GATAD2B were used as bait. Bait proteins were excluded from calculations, because they are in excess amount and introduce a bias in stoichiometry calculations. Stoichiometry of the subunits where **A)** MBD2/3 and **B)** GATAD2B were used as bait. **C)** Scatter plot show log2 of iBAQ intensities for HDAC1 and HDAC2 in all six replicates of CDK2AP1 PD-MS. **D)** Schematic representation of the possible new sub-complexes independent of the NuRD complex that can be formed via PPIs occurring with bridging subunits.

Approximately the same number of unique peptides were detected for HDAC1/2 (∼20) in NuRD subunits PD-MS, but no iBAQ intensity values was calculated for HDAC1 in CDK2AP1 PD-MS suggesting that only HDAC2 but not HDAC1 was co-purified with the CDK2AP1/NuRD complex (**Fig. 6C**). To ensure that this observation is not due to lack of HDAC1 expression issue, we further analysed the transcript and protein expression levels of both HDAC1/2 (RNA-Seq dataset from Human Protein Atlas [36] and shotgun proteomics dataset from ProteomicsDB [37]) in HEK293 cells. These data show that HDAC1/2 express at the same level, and thus absence of HDAC1 in CDK2AP1 pulldowns was not related to expression. We conclude that this observation is linked to the assembly and architecture and consequently interaction of the HDAC1 with the CDK2AP1/NuRD complex (**Supplemental Fig. 4A & B)**.

## Discussion

The MGCC module of the NuRD complex plays a diverse role in almost all stages of development and in many disease states in metazoans [38,39]. A central question concerning multi-subunit assemblies such as MGCC is how specificity is achieved whilst the complex is recruited to specific genomic loci. It is generally acknowledged that subunit-specific protein interactions provide some of this regulatory specificity to multi-protein complexes. For example, two groups recently demonstrated that PWWP2A, an H2A.Z binding protein, binds to MTA1 and separates MHR from MGCC, and thus forms PWWP2A-MTA1-HDAC1/2-RBBP4/7 complexes [40,41]. It is therefore plausible that NuRD subunit-specific binding partners that we report in this study could give rise to a combination of additional NuRD subcomplexes that would add to this functional diversity by creating other NuRD species with distinct compositions and stoichiometries. Our interactome data lay the foundation for future studies to investigate the functional readout of such complexes.

Recently, Sher et al. demonstrated that disruption of the MBD2/NuRD axis, but not MBD3/NuRD, leads to the derepression of fetal hemoglobin genes [15,16]. MBD2 is a methyl-CpG-binding protein; however, the proximal promoter of γ-globin and the entire β-globin locus is depleted of CpG islands, suggesting that MBD2 may not be acting to directly bind methylated DNA at the β-globin locus. Identification of a PPI network for MBD2a and MBD2b isoforms could potentially help to define a mechanism of their function and pave the way for more targeted molecular therapies. Here, we demonstrate that MBD2 isoforms co-purify with dozens of proteins involved in gene regulation and genome structure organisation; these proteins could potentially contribute to γ-globin repression. It is highly likely that other proteins – such as ZNF219 – might facilitate the binding of MBD2/NuRD to the β-globin gene locus.

Note that enrichment of ZNF219 and CDK2AP1 with MBD2 but not MBD3 does not necessarily indicate that they do not interact with MBD3/NuRD. Regarding ZNF219, we have previously reported the enrichment of ZNF219 with MBD3/NuRD in NTERA-2 cells, where MBD2 is not expressed [6]. Together, these data might indicate that there might be a competition between MBD2/NuRD and MBD3/NuRD for binding to ZNF219. Thus, in the presence of MBD2 the interaction of ZNF219-MBD3/NuRD could occur at very low stoichiometry and below the detection limit of our LC-MS/MS method in this cell line (HEK293). Our data reveal that ZNF219/MBD2b-NuRD and CDK2AP1/MBD2(a/b)-NuRD are more abundant compared to the MBD3-NuRD complex in HEK293 cells, Future functional studies of these complexes in relevant cell lines such as Human Umbilical Cord Derived Erythroid Progenitor-2 (HUDEP) cells could help define their function.

MBD PD-MS iBAQ data show substoichiometric ratio for CDK2AP1 and CHD4, which is consistent with previous studies. This may also indicate that the majority of MBD/NuRD complexes lack CHD and CDK2AP1 proteins. If true, the MBD/NuRD complex that carries both deacetylation and remodeling activities might be less abundant compared to MBD/NuRD with only deacetylase activity. The increase in ratios of MBD, CHD4, and CDK2AP1 when GATAD2B is used as bait might also imply that the MBD-GATAD2-CDK2AP1-CHD assembly is present as an independent complex with remodeling activity.

In conclusion, we report several specific protein interactions for the subunits of the MGCC module and further report the presence of new sub-complexes independent of the NuRD complex such as GATAD2-CDK2AP1-CHD and CDK2AP1/NCOR complexes **(Fig. 6D)**. Future studies may shed more light on the biological function of these interactors and sub-complexes in gene regulation in normal and disease states. The generation of recombinant protein complexes or tandem purification of the complexes followed by biophysical analysis might reveal the precise stoichiometry and structure and molecular function of these derivative complexes.

## Materials and Methods

### Plasmid Constructs

All genes used in this study were cloned into pcDNA3.1(+) vector using Gibson Assembly and were either N-terminally FLAG- or HA-tagged. Except for ADNP (obtained as a cDNA from Horizon Discovery (Genbank #BC075794)) and GATAD2B-C^Del^ constructs, the rest were a kind gift from Professor Joel Mackay, The University of Sydney. A list of primers used for Gibson assembly of all cDNAs into pcDNA3.1(+) are available on request.

### Design of APR peptides

The GATAD2A/B protein sequences were analysed by TANGO to detect aggregation prone regions [31]. Default physicochemical parameters were selected as below: temperature, 298 K; pH 7.5; ionic strength, 0.02 M; and concentration, 1 M. An aggregation score of 5 was set as a cut-off per residue. The residues spanning each aggregation prone region were combined with an 11 aa portion of HIV-1 Tat protein (YGRKKRRQRRR) to enhance cell permeability. Peptides were synthesized to at least 80% purity by HPLC at Mimotopes, Australia.

### Cell culture and transfection

K562 cells were grown in RPMI 1640 supplemented with 10% (v/v) foetal calf serum, penicillin (100 U/mL) and streptomycin (100 μg/mL). Expi293F™ cells were grown to a density of 1.5 × 10^6^ cells/mL in Expi293™ Expression Medium (Thermo Fisher Scientific). Combinations of equimolar quantities of constructs were co-transfected into cells using linear polyethylenimine (PEI) (Polysciences, Warrington, PA, USA). DNA (4 μg) was first diluted in 200 μL of PBS and vortexed briefly. PEI (8 uL, 1 mg/mL) was then added and the mixture was vortexed again, incubated for 20 min at room temperature, and then added to 1.9 mL of cells in a 12-well plate. The cells were incubated for 65-72 h at 37 °C, 5% CO2 in humidified incubator on a horizontal orbital shaker (130 rpm). Aliquots of cells (1 mL) were then harvested, washed twice with PBS, centrifuged (300 g, 5 min), snap-frozen in liquid nitrogen and stored at −80 °C.

### Cell lysate and APR treatment for aggregation analysis

K562 cells were lysed using 50 mM Tris-HCl, 150 mM NaCl, 0.5% IGEPAL, pH 7.5 and protease inhibitor cocktail (Sigma Aldrich), 1 mM DTT, and 1 μL Pierce™ Universal Nuclease (Thermo Fisher Scientific). Then cells were sonicated for 5 cycles, 1 min ON/10 s OFF. After sonication total lysate was collected and the rest of the lysate was aliquoted into new 1.5 mL Eppendorf tubes and then known concentration of APR peptides were added to each tube and incubated for 30 min at 4 °C. Next, tubes were spun at 20,000 *g* for 30 min to separate the soluble and insoluble fractions.

### *In vitro* protein expression and Co-IP

DNA constructs were transcribed and translated *in vitro* in pairs using equimolar plasmids in 70 μL of TNT® Quick Coupled Transcription/Translation System (Promega). RNaseOUT (0.5 ΜL, Thermo Fisher Scientific), methionine (2 μL, 1 mM) and 1x protease inhibitor (Sigma Aldrich) were added to each reaction. The reactions were incubated at 30 °C for 3 h. Prior to immunoprecipitation, 500 μL lysis buffer (50 mM Tris-HCl, 150 mM NaCl, 0.1% (v/v) Triton X-100, 1 mM DTT, pH 7.5) was added to the reactions. Input (5% of total) was collected and the remainder was mixed with 20 μL of anti-FLAG Sepharose 4B beads (Sigma Aldrich) at 4 °C for 2 h on a rotator. Beads then were washed with 500 μL of wash buffer (50 mM Tris-HCl, 200 or 500 mM NaCl, 0.5% (v/v) IGEPAL^®^ CA630, 0.2 mM DTT, pH 7.5) for 5 times. Finally, elution was performed three times each using 20 μL of a 150 μg/μL stock of 3xFLAG peptides (Sigma Aldrich). Western blot analysis was done as previously described [7]. Antibodies used in this study are as follows; GATAD2A (#A302-358A, Bethyl lab), GATAD2B (#A301-281A, Bethyl lab), FLAG (#A8592-1, Sigma Aldrich), HA (#2999, Cell Signalling Technology), and GAPDH (#ab8245, Abcam).

### Sample preparation and tandem mass spectrometry

Label-free FLAG pulldowns were performed in at least triplicate. Nuclear extracts of transiently transfected Expi293F cells were incubated with 20 μL anti-FLAG beads (Sigma Aldrich) After incubation for 2 h, five washes were performed: 3 washes with a buffer containing (200 mM NaCl, 50 mM Tris-HCl, 0.5% (v/v) IGEPAL, PH 7.5), and two washes with the same buffer lacking IGEPAL. Affinity-purified proteins were subject to on-bead trypsin digestion, where 20 μL of digestion buffer (2 M urea freshly dissolved in 50 mM Tris-HCl, 1 mM DTT, 100 ng trypsin, 20 ng LysC (Promega)) was added and then vortexed in 30 °C for 2 h. Next, the beads were collected, and supernatant was transferred into Lobind tubes. Beads were resuspended in 20 μL 2 M urea containing 10 mM IAA in the dark for 20 min. The supernatant was transferred to the previous tube and incubated at 30 °C for 16 h. The following day, tryptic peptides were acidified to a final concentration of 2% (v/v) with formic acid (Sigma Aldrich) and desalted using StageTips (Thermo Fisher Scientific). Peptides were dried in a SpeedyVac and dissolved in 10 μL 0.1% (v/v) formic acid. LC-MS/MS analysis was performed on an UltiMate™ 3000 RSLCnano System (Thermo Fisher Scientific) system connected to a Thermo Scientific Q-Exactive HF-X hybrid quadrupole-Orbitrap mass spectrometer equipped with a standard nano electrospray source (Thermo Fisher Scientific). Peptides (3 ΜL) were injected onto a C18 column (35 cm x 75 μm inner diameter column packed in-house with 1.9 μm C18AQ particles). Peptides were separated at a flow rate of 200 nL/min using a linear gradient of 5–30% buffer B over 30 min. Solvent A consisted of 0.1% (v/v) formic acid, and solvent B consisted of 80% (v/v) acetonitrile, 0.1% (v/v) formic acid. The end-to-end run time was 45 min, including sample loading and column equilibration times. The mass spectrometer was set to a data-dependent acquisition mode (DDA). In the DDA run, each full scan MS1 was operated as follows: mass scan range was between 300-1600 m/z at resolution of 60,000. The top 15 most intense precursor ions were selected to be fragmented in the Orbitrap via high-energy collision dissociation activation. MS2 scan was operated as follow; mass scan range was between 200-2000 m/z at resolution of 15,000, 1 × 10^5^ AGC target, and 1.4 m/z isolation window.

### Raw data analysis

Raw data were analysed by MaxQuant (version 1.6.6.0) [44] using standard settings. Additional parameters including carbamidomethyl cysteine (C) and methionine oxidation (M) were selected as fixed and variable modifications, respectively, and LysC, Trypsin, were selected as proteolytic enzymes. The human proteome (Proteome ID UP000005640) was used as the reference proteome. The generated proteingroups.txt table in conjunction with an experimental design text file was used to perform all statistical analyses using LFQ values in R studio as described elsewhere [45]. Perseus algorithm was used to impute the missing values. Proteins that had two missing values were discarded from the analysis [46]. For better data visualisation the output files were further processed. First, all proteins with fold change >2 and statistically significant were kept. Then, proteins including heat shock, ribosomal, keratin or proteins with mitochondrial and cytoplasmic localisation were excluded from the list. A full list of unfiltered interactors is listed in the **Supplemental Table 1, S1**. To determine the stoichiometry of the subunits, the iBAQ-adjusted intensity values (iBAQ; is an approximate calculation of protein copy numbers by dividing the sum of intensities of all experimentally detected peptides by the number of theoretically observable peptides for each protein) for MTAs were averaged and then intensity values for known NuRD components were divided by the MTA value and multiplied by 2 because based on published X-ray crystal structure two molecules of HDAC1 and two molecules of MTA1 make the core of the NuRD complex [18]. Due to high sequence similarity between paralogues (i.e., RBBP4 is 92% identical to RBBP7) we first considered each set of paralogues as a single group. NetworkAnalyst tool was used for network visualisation of the unique and shared interactors. Of note, LFQ and iBAQ are two different quantification algorithms; iBAQ is an approximate calculation of protein copy numbers and is the best at determining ratio changes within samples not across samples. Whereas LFQ is the best representative of ratio changes between samples.

## Acknowledgments

We thank Cynthia Metierre for assisting with MTT assays. We also appreciate the technical assistance we received from the Mass Spectrometry core facility at The University of Sydney.

## Author Contributions

M.S.T and C.G.B conceived the study, designed MS experiments, analysed data and wrote the manuscript. C.G., J.F., and H.F. prepared samples for MS experiments and performed western blots and MMT assays. J.K.K.L. designed and cloned constructs and reviewed the manuscript. J.P.M. reviewed the manuscript and provided intellectual input on the studies and manuscript. J.E.J.R. reviewed the manuscript and provided intellectual input, research governance and scientific discussion.

## Supplemental Figure Legends

**Supplemental Fig. 1**. Bar graph indicates that CDK2AP1 interacts with GATA2B-C (C-terminal segment) as compared to GATADB-N (N-terminal region). No iBAQ value was calculated for CDK2AP1 protein when GATAD2B-N was used as bait.

**Supplemental Fig. 2**. Volcano plot showing the log2 fold change plotted against log10 adjusted P value for the middle section of CHD4 (CHD4-M) PD-MS. Most of the enriched proteins are known to play a role in chromatin biology.

**Supplemental Fig. 3**. Cancer-associated missense mutation in the C-terminal region of CHD4 may change the balance of CHD4/NuRD vs CHD4/ChAHP. A FLAG-tagged GATAD2B or ADNP were coexpressed with HA-CHD4-C [wild-type (WT) or mutants] by *in vitro* transcription–translation (IVT) in a rabbit reticulocyte lysate. **A)** list of mutations examined in the pairwise interaction experiments. **B)** FLAG-fusion proteins were immobilized on FLAG affinity beads and used as baits to pull down the coexpressed HA-CHD4-C. FLAG peptides were used to elute off the FLAG-tagged baits and interactors.

**Supplemental Fig. 4**. HDAC1 and HDAC2 exhibit equal abundance at both the mRNA and protein level in HEK293 cells. **A)** Normalised relative mRNA expression for HDAC1 and HDAC2 reported in Human Protein Atlas [36]. **B)** Normalised protein expression of HDAC1 and HDAC2, the normalised iBAQ value is reported in ProteomicsDB [37].

**Supplemental Table 2**. Number of unique peptides detected for each subunit of the NCOR complex and their representation in the Contaminant Repository for Affinity Purification Mass Spectrometry Data (CRAPome) [43].

## Data availability

The data that support the findings of this study are available from the corresponding author on request.

