## Supplementary figures and images for "Unique Protein Interaction Networks Define The Chromatin Remodeling Module of The NuRD Complex"

### Supplemental Fig1

#### Supplemental Figure 1

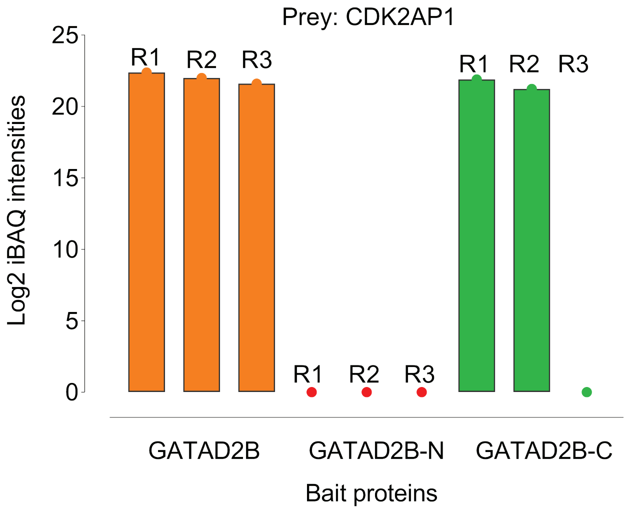

### Supplemental Fig2

#### Supplemental Figure 2

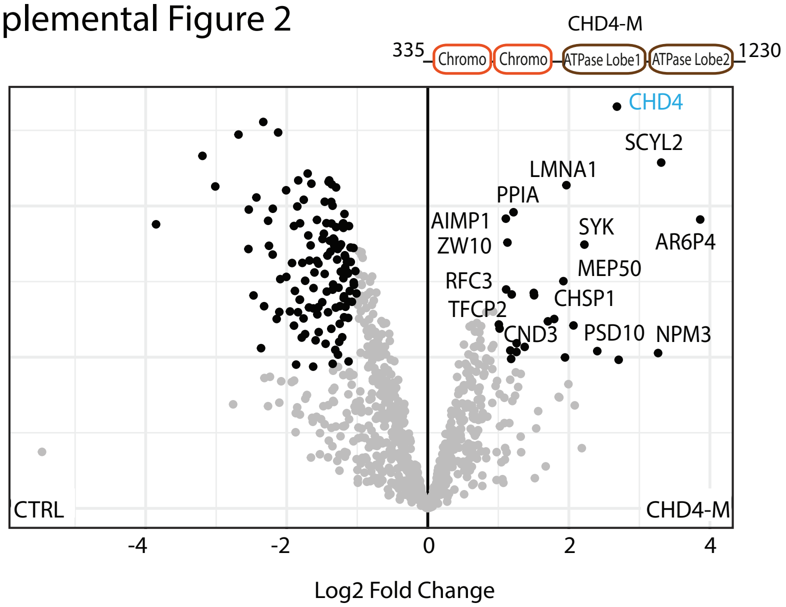

### Supplemental Fig4

Supplemental Figure 4

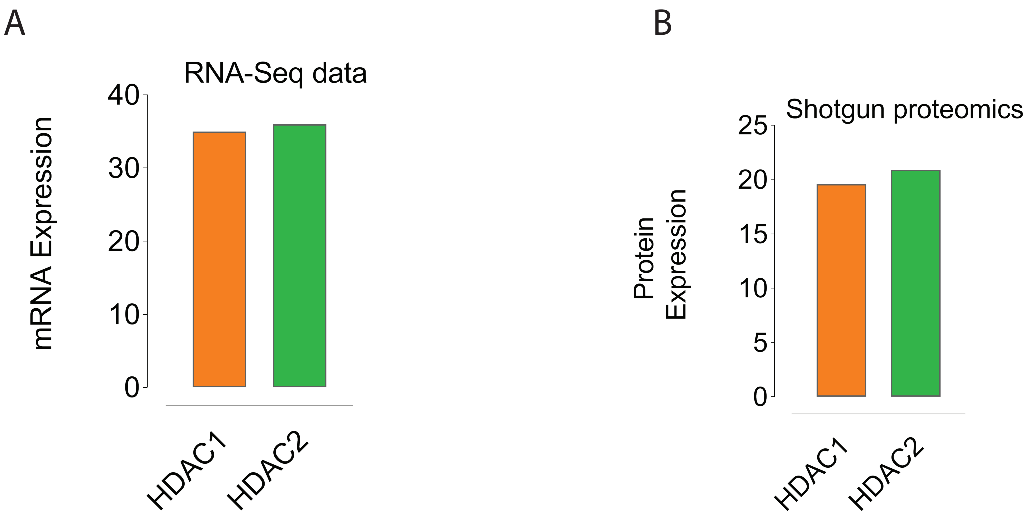
