## Supplemental Fig3 for "Unique Protein Interaction Networks Define The Chromatin Remodeling Module of The NuRD Complex"

### Supplemental Figure 3

A

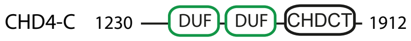


| Mutation | | | Cancer Type |
|---|---|---|---|
| A | 1866 | D | Uterine Endometrioid Carcinoma |
| D | 1867 | N | Breast Invasive Ductal Carcinoma |
| P | 1879 | S | Cutaneous Melanoma |
| E | 1889 | K | Lung Adenocarcinoma |
| R | 1890 | C | Colorectal Adenocarcinoma |
| N | 1891 | D | Stomach Adenocarcinoma |

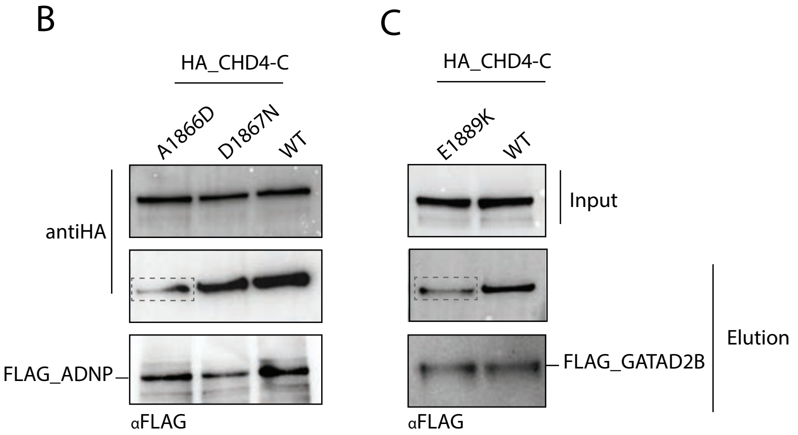
