## Supplemental Table 2 for "Unique Protein Interaction Networks Define The Chromatin Remodeling Module of The NuRD Complex"

| Protein | No. Unique Peptides | CRAPome Score | Localisation |
|---|---|---|---|
| HDAC3 | 6 | 8 | Nucleus |
| NCOR1 | 69 | 23 | Nucleus |
| NCOR2 | 56 | 14 | Nucleus |
| TBL1R | 11 | 0 | Nucleus |
| GPS2 | 9 | 18 | Nucleus |
| TBL1X | 6 | 3 | Nucleus |
